# Beta secretase 1-dependent amyloid precursor protein processing promotes excessive vascular sprouting through NOTCH3 signaling

**DOI:** 10.1101/585489

**Authors:** Claire S. Durrant, Karsten Ruscher, Olivia Sheppard, Michael P. Coleman, Ilknur Özen

## Abstract

Amyloid beta peptides (Aβ) proteins play a key role in vascular pathology in Alzheimer’s Disease (AD) including impairment of the blood brain barrier and aberrant angiogenesis. Although previous work has demonstrated a pro-angiogenic role of Aβ, the exact mechanisms by which amyloid precursor protein (APP) processing and endothelial angiogenic signalling cascades interact in AD remain a largely unsolved problem. Here, we report that increased endothelial sprouting in human-APP transgenic mouse (TgCRND8) tissue is dependent on β-secretase (BACE1) processing of APP. Higher levels of Aβ processing in TgCRND8 tissue coincides with decreased NOTCH3/JAG1 signalling, over-production of endothelial filopodia and increased numbers of vascular pericytes. Using a novel *in vitro* approach to study sprouting angiogenesis in TgCRND8 organotypic brain slice cultures (OBSCs), we find that BACE1 inhibition normalises excessive endothelial filopodia formation and restores NOTCH3 signalling. These data present the first evidence for the potential of BACE1 inhibition as an effective therapeutic target for aberrant angiogenesis in AD.

**Significance:** In this study, we show that targeting amyloid beta processing provides an opportunity to selectively target tip cell filopodia-driven angiogenesis and develop therapeutic targets for vascular dysfunction related to aberrant angiogenesis in AD. Our data provide the first evidence for a safe level of BACE1 inhibition that can normalize excess angiogenesis in AD, without inducing vascular deficits in healthy tissue. Our findings may pave the way for the development of new angiogenesis dependent therapeutic strategies in Alzheimer’s Disease.

## Introduction

Alzheimer’s disease (AD) is closely associated with alterations in the vascular system. Multiple studies in humans and animal models have described pathological vascular changes in AD (1), including disruption to the neurovascular unit (2) and blood-brain barrier (3), increased microvessel density (2, 4, 5), arteriolar and venular tortuosity (6, 7), vascular Aβ accumulation (8) and alterations in retinal vasculature (9). Such changes will likely compromise the effective delivery of oxygen and nutrients to the brain, so understanding whether vascular alterations are a cause or consequence of aspects of AD pathology, notably Aβ accumulation, is required in order to design effective therapies.

Amyloid beta peptides, particularly Aβ_1-42_, are hallmarks of AD (10). These peptides are the result of sequential proteolytic cleavage of amyloid precursor protein (APP) by β- and γ-secretase enzyme activity. Whilst it is reported that synapse loss is the best correlate of clinical outcome in AD (11), it is unclear as to whether pathological APP processing products drive this effect by a direct action on neurons, or indirectly such as through aberrant angiogenesis. Despite widespread interest in the role of brain vasculature in AD, little is known about how amyloid-induced vascular changes alter pathological sprouting angiogenesis.

Endothelial sprouting is a key process of physiological and pathological angiogenesis (12) and is regulated by the NOTCH signaling pathway (13). NOTCH receptors (NOTCH1-4) undergo proteolytic processing via γ-secretase in a manner comparable to that of APP (14, 15) resulting in the hypothesis that interactions between these signaling pathways could underlie the angiogenic pathology in AD (16, 17). Pericyte recruitment is closely linked to endothelial cell sprouting. Endothelial tip cells secrete platelet derived growth factor (PDGF) that activates platelet derived growth factor receptor beta (PDGFRβ) on pericytes and induce their migration to the sprout (18, 19).

Organotypic brain slice cultures (OBSCs) provide an excellent platform to explore mechanisms of pathological sprouting angiogenesis in AD. OBSCs initially retain a dense network of capillaries and neurovascular units alongside maintenance of neuronal architecture and non-neuronal cell populations (20–22). Crucially, OBSCs provide a 3-dimensional culture system that supports the formation of new blood vessels and are amenable to pharmacological manipulation, live imaging and repeated measurements (23–26). We and others have previously shown that OBSCs are powerful tools for investigating the progression of AD-like alterations including Aβ accumulation (27), synaptic disruption (24) and cerebrovascular damage (28).

In this study, we used a novel *in vitro* approach to study sprouting angiogenesis in OBSCs. In tissue from huAPP transgenic mice, we found an increase in sprouting angiogenesis that could be completely blocked by BACE1 inhibition. We investigate the mechanisms by which pathological APP processing and NOTCH signalling interact to induce excessive vascular sprouting and discuss the implications for the blood vessel pathology seen in AD.

## Results

### Characterisation of sprouting angiogenesis in organotypic brain slice cultures

New blood vessels in the cortex are formed via sprouting angiogenesis, a process driven by specific endothelial tip cells at the leading edge of vascular sprouting. Sprouting angiogenesis encompasses sequential events including; sprouting at the vascular front of endothelial cells, extension of sprouts, the formation of new vascular loops, and pericyte recruitment (12, 29, 30). We first established an *ex vivo* cortical organotypic brain slice culture (OBSC) system to analyse this highly regulated process and found we were able to identify multiple stages of sprouting angiogenesis in OBSCs from wild-type (WT) mice (summarised in **Fig. 1e**). PECAM-1 staining (a marker of endothelial cells) showed a dense network of blood vessels expressing basal membrane protein laminin in 7-days *in vitro* cortical slices (**Fig. 1a**). Endothelial tip cells were found either along the trunk of PECAM-1+ blood vessels with few filopodia (**Fig. 1b, asterisk**) or at the leading edge of vascular sprouts extending numerous filopodia (**Fig. 1b, c**), indicating active sprouting angiogenesis. The filopodia of some vascular sprouts were also found to engage with those of a nearby endothelial tip cells to form a bridge and the formation of new blood vessels (**Fig. 1d**).

**Figure 1.**
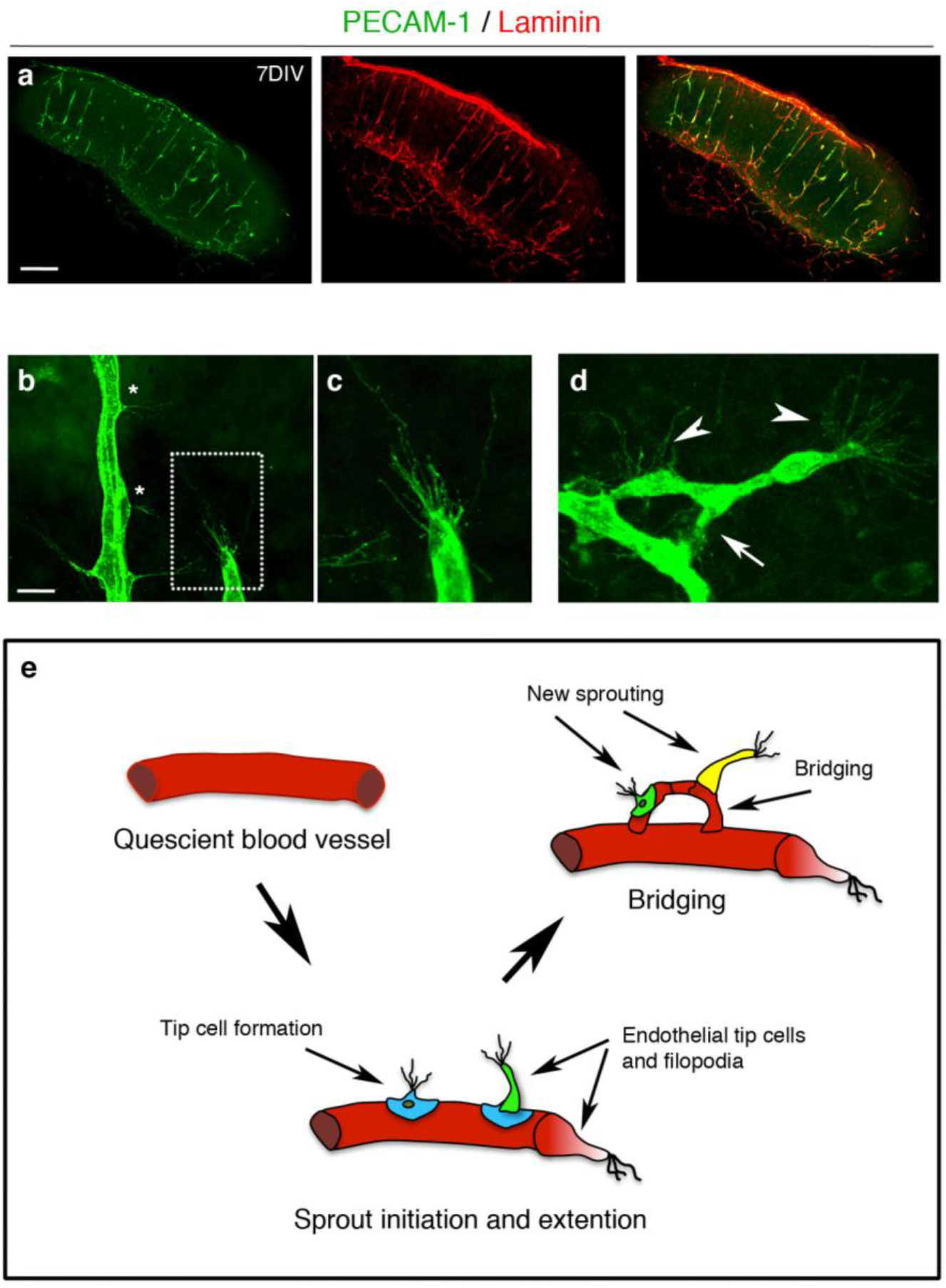
Characterisation of sprouting angiogenesis in 7-days *in vitro* organotypic cortical slice cultures from wildtype mice. **(a)** Confocal images of cortical slices stained for PECAM-1 (green) and laminin (red) to visualize blood vessels at 7 days *in vitro*, scale bar 500 μm **(b-d)** Different stages of vascular sprouting can be visualised in cortical brain slices including; tip cell formation (*) endothelial tip extension (framed area in **(b)** which is expanded in **(c)**) new sprouting (arrow heads) and bridging (arrow) scale bar 50 μm) **(d)**. **(e)** Diagram summarising the different stages of sprouting angiogenesis that can be observed in cortical organotypic slice cultures.

PDGFRβ^+^ pericytes were found around the OBSC blood vessels and with long cytoplasmic processes surrounding the abluminal surface of endothelium (**Fig. 2a-c**). High magnification confocal imaging showed that PDGFRβ expressing pericytes were closely associated with the PECAM-1+ vascular sprouts (**Fig. 2d-e**, framed image) and astrocytes within neurovascular unit (**Fig. 2f-h**) at 7 days *in vitro*. Taken together, this demonstrates the utility of OBSCs as a tool to assess different steps of sprouting angiogenesis, including filopodia formation and pericyte coverage.

**Figure 2.**
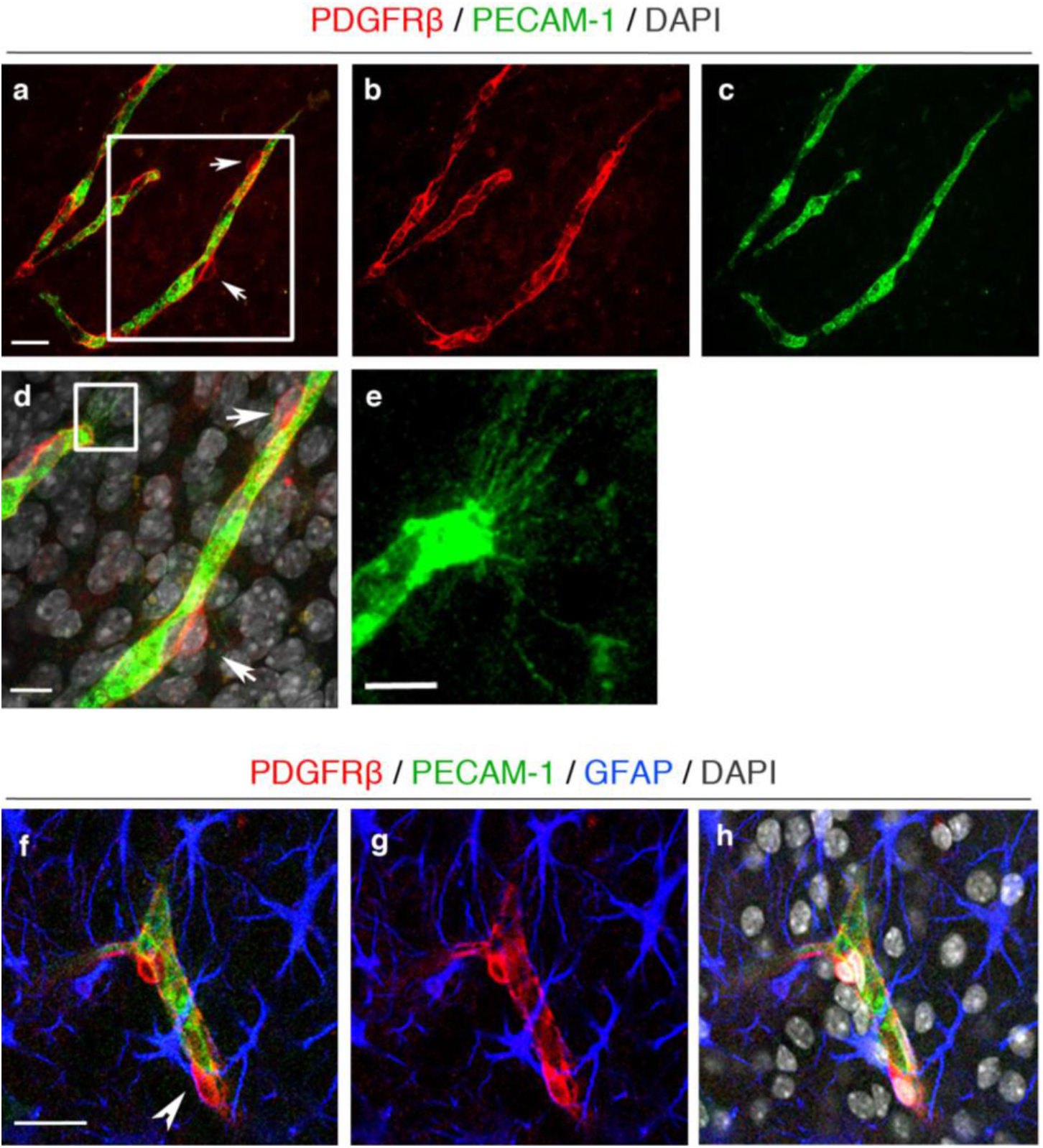
Pericytes are well preserved in organotypic cortical slices at 7 days *in vitro*. **(a-c)** Confocal images showing PECAM-1+ blood vessels (green) **(a, c)** covered by PDGFRβ+ pericytes (red, arrows **(a, b)**) in 7 days *in vitro* WT cortical slices; scale bar 20 μm. **(d)** Higher magnification images of the framed area in **(a)** with DAPI counterstain depicting nuclei (grey). Pericytes (arrows) can be seen on the PECAM-1+ angiogenic blood vessels; scale bar 10 μm. **(e)** depicts enlarged area framed in **(d)** showing clear vascular sprouting. **(f-h)** Confocal z-stacks show a preserved cytoarchitecture, consisting of PECAM-1+ blood vessels (green), surrounded by PDGFRβ^+^ pericytes (red) and GFAP+ astrocytes (blue) (DAPI=grey). Arrow marks the close association between the pericytes and astrocytes; scale bar 20 μm.

### Vascular abnormalities in the cortex and retina of postnatal TgCRND8 mice

In preparation for studying the mechanism in OBSCs, which necessarily require the use of early postnatal tissue, we studied P7 TgCRND8 post mortem mouse cortex and retina to determine whether a mutant huAPP transgene leads to any pathological changes in the organisation of the vascular network and endothelial cell physiology during postnatal development (31, 32). We first analysed vessel architecture and pericyte coverage in the cortex of P7 mice (**Fig. 3**). PECAM-1 expressing cortical blood vessels of P7 TgCRND8 (**Fig. 3b, b’**) mice appeared to be more tortuous than cortical vessels of P7 WT mice (**Fig. 3a, a’**). When assessing PDGFRβ+ pericyte coverage of blood vessels in P7 TgCRND8 cortex, we found an approximately 2-fold higher pericyte coverage when compared to WT (**Fig. 3c**) with coverage becoming almost complete (**Fig. 3b**). We next analysed the vascular organisation of the retinal plexus in P7 WT and TgCRND8 animals (**Fig. 3d**). PECAM-1+ blood vessels in the retinal plexus of TgCRND8 were larger in diameter and formed a highly dense and more interconnected network compared to WT littermates, indicating aberrant angiogenesis. Thus, several CNS tissues from mutant huAPP transgenic mice show enhanced angiogenesis as early as P7.

**Figure 3.**
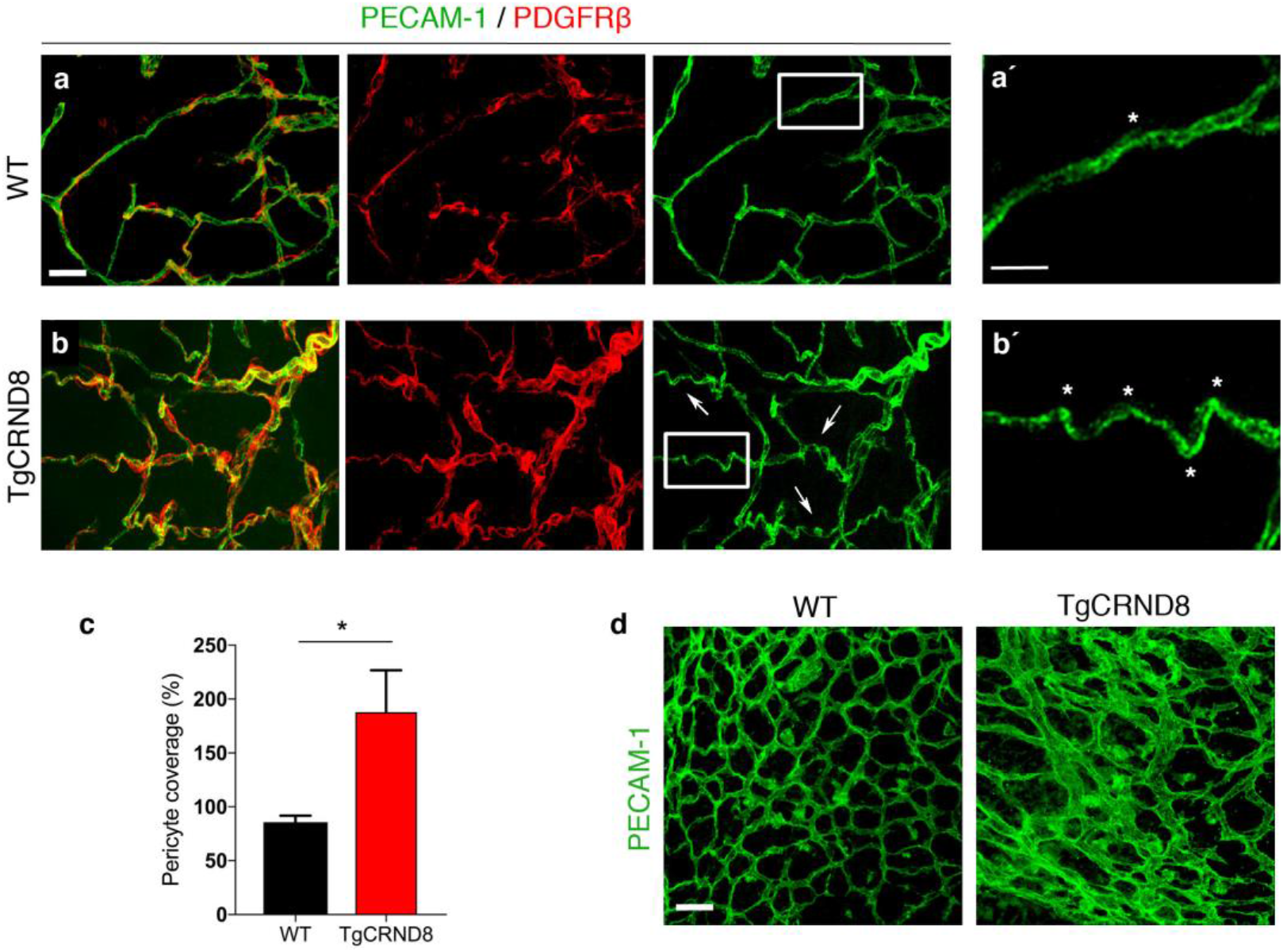
Vascular morphology in the cortex and retina of postnatal TgCRND8 mice. **(a-b)** Blood vessels were stained by PECAM-1 and pericytes by PDGFRβ. Increased pericyte coverage and microvessel tortuosity (arrows and framed areas) is seen in P7 TgCRND8 cortex; scale bar 20 μm. **(a’-b’)** Enlarged white-framed areas in a and b showing samples of normal microvessels in WT **(a’)** and tortuous microvessels (*) in TgCRDN8 **(b’)** cortex; scale bar 10 μm. **(c)** There is increased PDGFRβ+ pericyte coverage in TgCRND8 cortex; mean ± SD (n=2 (WT), n=4 (TgCRND8), *P<0.05 Student’s t-test. **(d)** Representative confocal images of PECAM-1 stained flat retinal plexus in P7 WT and TgCRDN8 mice; scale bar 20 μm.

### Excessive vascular sprouting is associated with increased vascular density and number of pericytes in TgCRND8 OBSCs

We next looked for changes in endothelial filopodia formation, vascular density and pericyte number in 7 days *in vitro* TgCRND8 OBSCs (**Fig. 4**). The capillary density (as measured by area of PECAM-1 staining) was significantly higher in TgCRND8 OBSCs compared to cultures from WT littermates (**Fig. 4a-b**). Confocal microscopy revealed an increase in the number of filopodia found both at the leading edge, as well as along the body, of the vascular sprouts in TgCRND8 cultures (**Fig. 4c**) alongside a 50% increase in the number of filopodia at the vascular sprout of the blood vessels (**Fig. 4d**).

**Figure 4.**
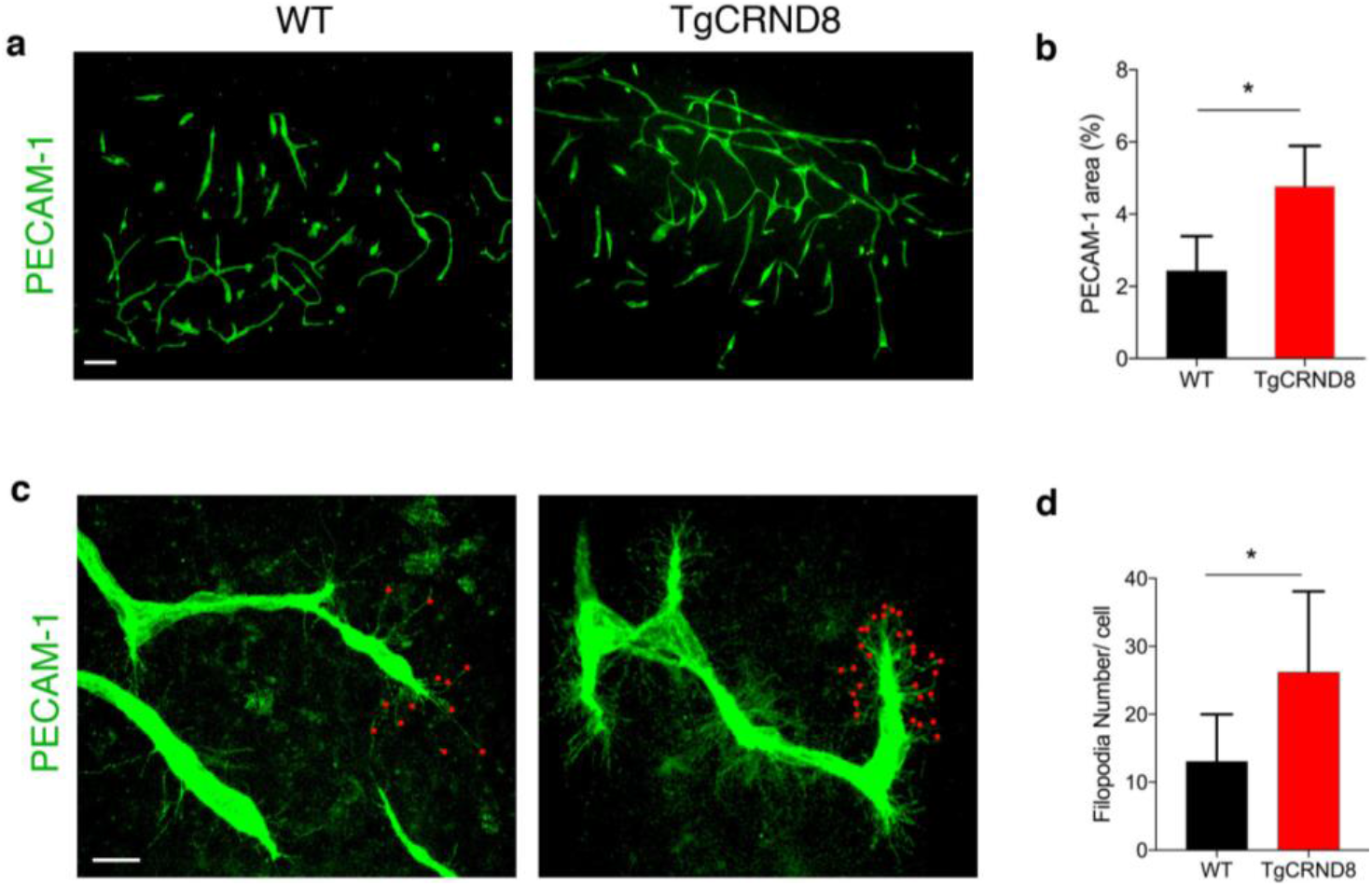
TgCRND8 organotypic cortical slices show increased vascular density and excessive filopodia formation when compared to wildtype cultures. **(a)** Representative confocal images showing blood vessel density (PECAM-1, green) in 7 days *in vitro* WT and TgCRDN8 slices; scale bar 100 μm. **(b)** Quantification of PECAM-1+ area (as % of the total image) reveals a significantly higher blood vessel density in TgCRND8 cortical slices versus WT slices. (mean ± SD (n=5 (WT), n=4 (CRND8), *P<0.05 Student’s t-test.) **(c)** Confocal images showing endothelial cells extending numerous finger-like filopodia at the forefront of vascular sprouts in 7 days *in vitro* WT and TgCRND8 cortical slices, visualized by PECAM-1 labelling. Red dots highlight vascular sprouting tips, scale bar 20μm. **(d)** Quantification shows that the number of filopodia per cell is significantly higher in TgCRND8 when compared to WT slices. (mean ± SD, (n=14 (WT), n=13 (TgCRND8), *P<0.05 Student’s t-test)

In 7 days *in vitro* TgCRND8 OBSCs, the number of PDGFRβ+ pericytes around the capillaries was significantly higher than WT controls (**Fig. 5a-b**). The increased number of PDGFRβ+ pericytes was correlated with an upregulation of PDGFRβ protein expression as measured by western blot (**Fig. 5c-d**).

**Figure 5.**
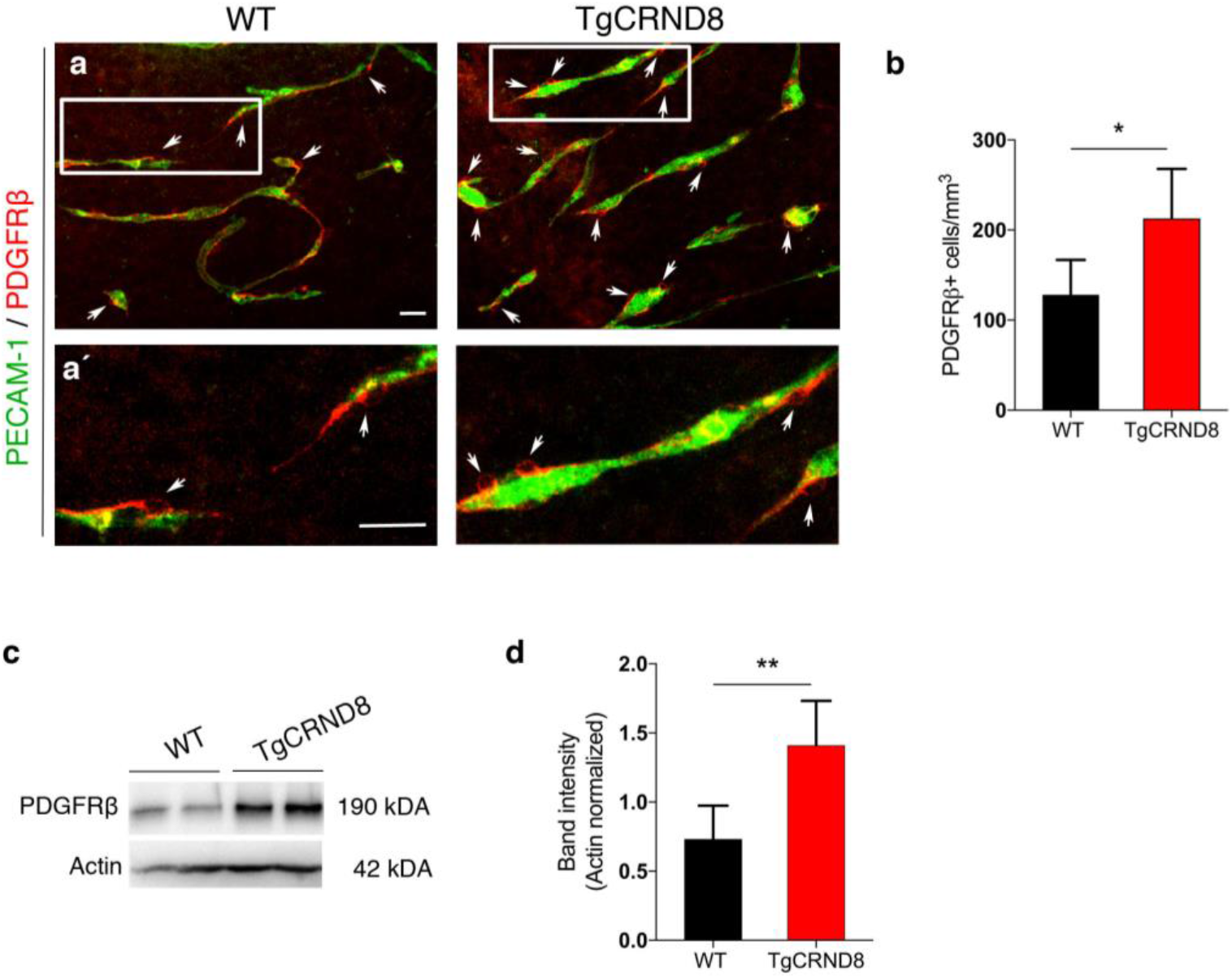
Increased pericyte number and PDGFRβ expression in TgCRND8 organotypic cortical slices. **(a)** Representative confocal images showing PDGFRβ+ pericytes (arrows) around blood vessels (PECAM-1, green) in 7 days *in vitro* WT and TgCRND8 slices; scale bar 50 μm. PDGFRβ+ pericytes are closely associated with cortical microvessels. **(a’)** The framed areas in **(a)** are enlarged, scale bar 50 μm **(b)** Quantification of PDGFRβ+ pericytes reveals an increased number in 7 days *in vitro* TgCRND8 slices when compared to WT (data are expressed as cell numbers per mm^3^; mean ± SD. (n=4 (WT), n=5 (TgCRND8), *P<0.05 Student’s t-test). Representative Western blots **(c)** and quantification of PDGFRβ band intensity **(d)** in 7 days *in vitro* WT and TgCRND8 cortical slices, shows increased PDGFRβ in TgCRND8 cultures when normalised to Actin control (n=5 (WT), n=4 (TgCRDN8) **P<0.01 Student’s t-test).

### Inhibition of BACE1 activity normalises vascular density and hypersprouting in TgCRND8 OBSCs

To investigate if higher vascular density and filopodia number depend on the increased production of Aβ seen in TgCRND8 OBSCs, we applied the BACE1 inhibitor LY2886721 to OBSCs for 7 days *in vitro* (**Fig. 6a**). Using ELISA, we found that BACE1 inhibitor completely abolished the generation of Aβ_1-40_ and Aβ_1-42_ in the TgCRND8 OBSC culture medium (**Fig. 6b**). Staining for PECAM-1, we found that BACE1 inhibition reduced the vascular density in TgCRND8 slices back to WT levels, with no additional effect on WT cultures (**Fig. 6c-e**). Quantification of PECAM-1+ capillaries revealed a two-fold increase in total vessel length in TgCRND8 OBSCs, which was restored to WT levels after BACE1 inhibition (**Fig. 6f**). Similarly, BACE1 inhibition reduced the excessive sprouting activity of PECAM-1+ endothelial cells in TgCRND8 OBSCs, significantly lowering the number of filopodia at the leading edge of the vascular sprout back to WT levels (**Fig. 6g-i**).

**Figure 6.**
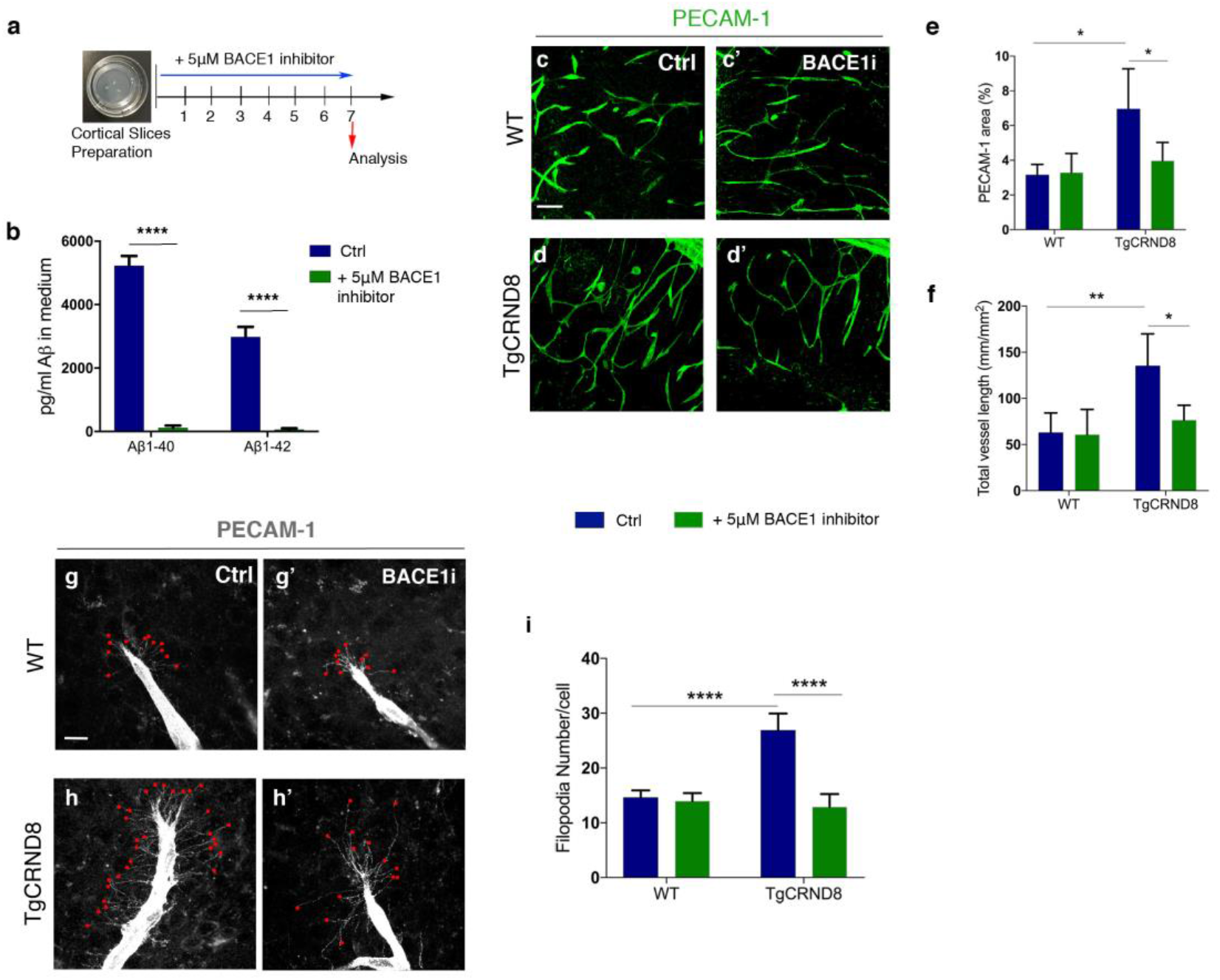
BACE1 inhibition decreases vascular density and normalises aberrant vascular sprouting in TgCRND8 organotypic cortical slices. **(a)** Diagram showing the experimental schedule for BACE1 inhibitor treatment of WT and TgCRND8 cortical slices. **(b)** Measurement of Aβ_1-40_ and Aβ_1-42_ in the culture medium of 7 day *in vitro* TgCRND8 slices treated with BACE1 inhibitor (mean ± SEM (n=4), two way ANOVA effect of treatment ****P<0.0001). **(c)** Confocal images showing blood vessel density (PECAM-1, green) in 7 days *in vitro* control **(c)** and BACE1 inhibitor **(c’)** treated WT slices. **(d-d’)** Confocal images showing blood vessel density (PECAM-1, green) in 7 days *in vitro* control **(d)** and BACE1 inhibitor **(d’)** treated TgCRND8 slices; scale bar 20 μm. BACE1 inhibition rescues the increase in PECAM1+ area (% image coverage) (mean ± SD (n=4 (WT), n=4 (TgCRND8), *P<0.05, two-way ANOVA) **(e)** and total vessel length (mm/mm^2^) (mean ± SD (n=4 (WT), n=4 (TgCRDN8), **P<0.01 and *P<0.05, two-way ANOVA) **(f)** seen in control treated TgCRND8 cultures. PECAM-1 staining of WT **(g-g’)** and TgCRND8 **(h-h’)** slice cultures with **(g’, h’)** or without **(g, h)** BACE1 inhibitor treatment. Individual filopodia are highlighted with red dots. **(i)** Quantification of the number of filopodia per cell demonstrates that BACE1 inhibitor rescues the increased number seen in untreated TgCRND8 cultures when compared to WT at 7 days *in vitro* (mean ± SD, (n=4 (WT), n=4 (TgCRND8), ****P<0.0001, two-way ANOVA).

### BACE1 inhibition restores *Notch3* and *Jag1* mRNA levels in TgCRND8 cortical slices

Given that modulating APP/Aβ metabolism via BACE1 inhibition resulted in normalisation of hypersprouting, we hypothesised that interaction between Aβ peptide processing and NOTCH signalling might explain the hypersprouting of the blood vessels observed in TgCRND8 mice. To test this hypothesis, we first examined the mRNA levels of key components of the NOTCH signalling pathway, NOTCH1, NOTCH3, JAG1, JAG2 and *DLL4*, in retinal tissue from postnatal TgCRND8 mice. Real time quantitative PCR analysis showed that mRNA levels of *Notch3* and *Jag1* were significantly lower in TgCRND8 retina when compared to the WT controls, whilst expression of *Notch1, Jag2* and *DLL4* were not significantly changed (**Fig. 7a**). Next, we asked whether BACE1 inhibition could rescue mRNA levels of *Notch3* and *Jag1* in TgCRND8 OBSCs. 5μM BACEI inhibitor treatment for 7 days *in vitro* normalised both *Notch3* and *Jag1* mRNA expression back to the levels observed in WT cultures (**Fig. 7b, c**). We found no significant changes in the mRNA expression of *Notch1, Jag2, DLL4* in TgCRND8 or WT slices after BACE1 inhibitor treatment (**Fig. 7d-e**).

**Figure 7.**
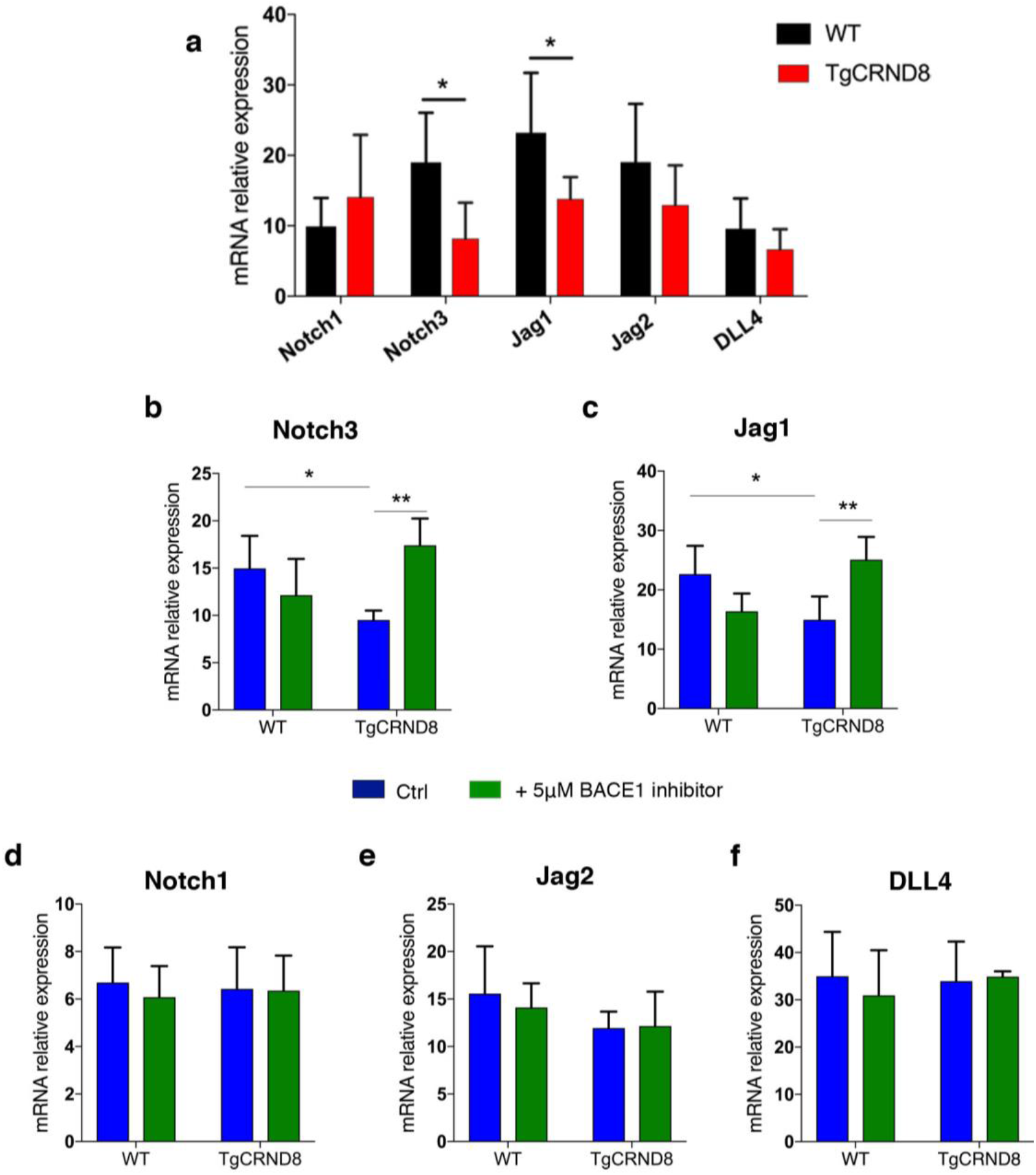
NOTCH signalling related gene expression in P7 retina and BACE1-treated organotypic cortical slices. **(a)** Quantitative gene expression analysis of NOTCH receptors (*Notch1* and *Notch3*) and NOTCH ligands (*Dll4, Jag1, Jag2*) in the retina of P7 WT and TgCRDN8 mice (mean ± SD, n=7 (WT), n=6 (TgCRND8), *P<0.05 Student’s t-test.) **(b-f)** Quantitative gene expression analysis of NOTCH receptors (*Notch1* and -*3*) and NOTCH ligands (*Dll4, Jag1, Jag2*) in 7 days *in vitro* WT and TgCRND8 cortical slices treated with BACE1 inhibitor. BACE1 inhibitor treatment normalised the expression levels of *Notch3* **(b)** and *Jag1* **(c)** in TgCRND8 cortical slices (mean ± SD, n=5 (WT), n=6 (TgCRDN8), *P<0.05 and **P<0.01, two-way ANOVA). BACE1 inhibitor treatment had no effect on the expression of *Notch1* **(d)**, *Jag2* **(e)**, and *DLL4* **(f)** in 7 days *in vitro* TgCRND8 or WT cortical slices, (mean ± SD, n=5 (WT), n=6 (TgCRND8), P>0.05, two-way ANOVA).

### BACE1 inhibition induces cleavage of NOTCH3 intracellular domain (NICD3) in TgCRND8 cortical slices

Finally, we analysed how higher production of Aβ affects NOTCH3 and NOTCH1 signalling activity in TgCRND8 OBSCs (**Fig. 8**). Translocation of the NOTCH intracellular domain (NICD) into the nucleus is a regulator of endothelial sprouting (13), so we tested whether the reduction in *Notch3* mRNA led to lower levels of NICD3. Western blot analysis showed a trend for reduced levels of NOTCH3 intracellular domain (NICD3) in TgCRND8 cortical slices. In contrast, BACE1 inhibitor treatment significantly increased NICD3 levels in TgCRND8 slices to at least the level of WT cortical cultures (**Fig. 8a-b**). Consistent with the mRNA levels of *Notch1*, there was no effect of genotype or BACE1 inhibition on the appearance of NOTCH1 intracellular domain (NICD1) (**Fig. 8c-d**).

**Figure 8.**
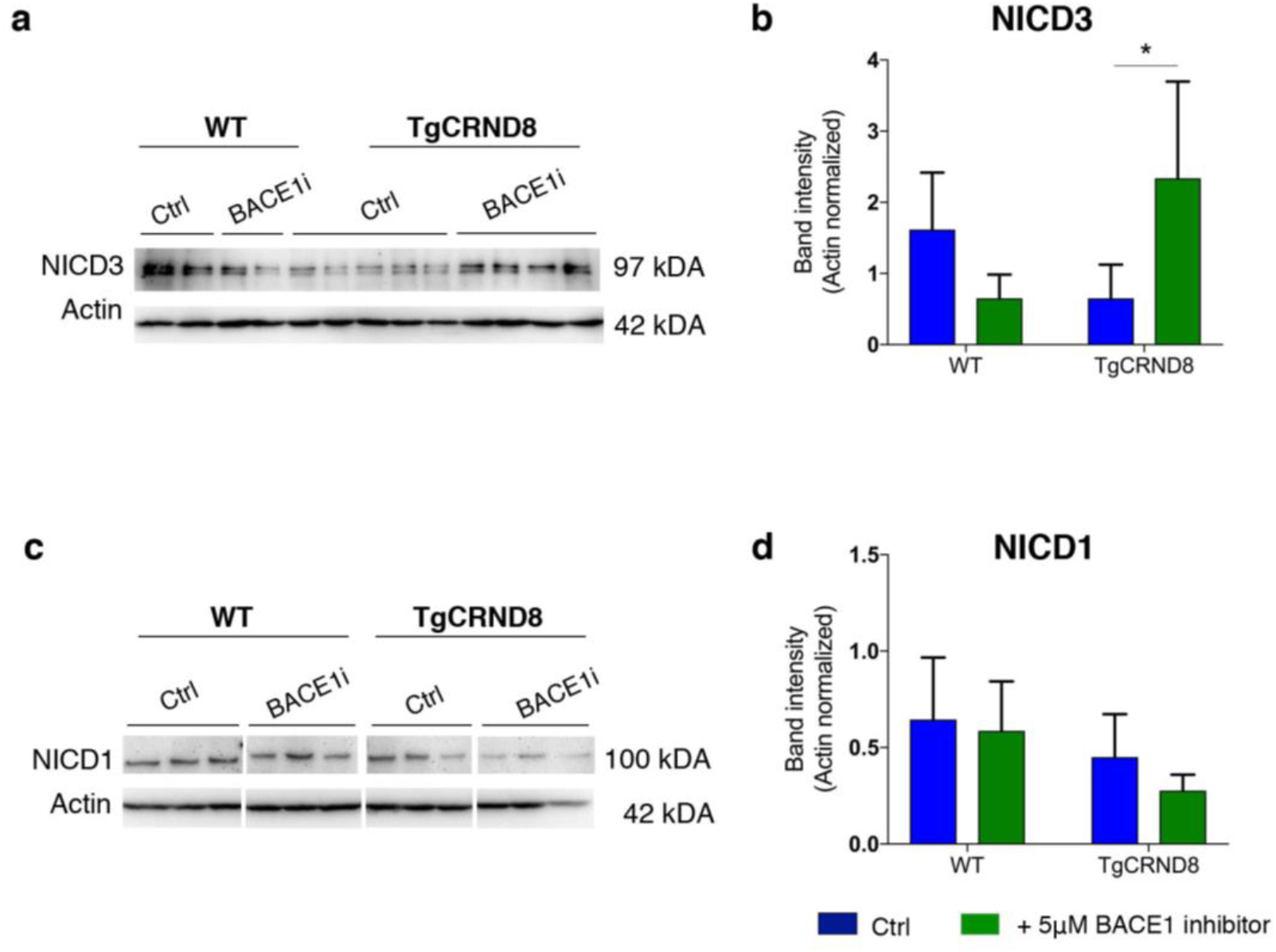
BACE1 inhibitor increases NOTCH3 intracellular domain (NICD3) in TgCRND8 organotypic cortical slices. **(a-b)** Representative Western blots and quantification of NOTCH3 intracellular domain (NICD3) in 7 days *in vitro* WT and TgCRND8 cortical slices treated with BACE1 inhibitor. (Data expressed in band intensity; mean ± SD, *P<0.05, n=4 (WT), n=12 (TgCRND8), two-way ANOVA, Tukey post hoc test.) **(c-d)** Representative Western blots and quantification of Notch1 intracellular domain (NICD1) in 7 days *in vitro* WT and TgCRND8 cortical slices treated with BACE1 inhibitor (Data expressed in band intensity; mean ± SD, n=9 (WT), n=14 (TgCRND8), P>0.05, two-way ANOVA).

## Discussion

In this study, we have shown that aspects of vascular pathology seen in AD can be recapitulated in an *in vitro* system that not only retains microvasculature but supports the growth of new vessels and is amenable to pharmacological manipulation. We show increased angiogenesis in postnatal TgCRND8 cortex, retina and in OBSCs. Using OBSCs, we demonstrated reversal of angiogenic pathology via BACE1 inhibition, alongside restoration of NOTCH signalling.

OBSCs represent a potent tool for exploring vascular phenotypes in a system that retains native cell populations and cytoarchitecture, Until now, whilst the OBSC model has been well established for studying angiogenic processes, (33), (34), (20), there were few studies seeking to explore this in combination with AD models. Slices from adult APP_SweDI mice have been found to express L-type calcium channels and substance P around amyloid plaques, pointing to altered angiogenesis at this key pathological site (35). Another study demonstrated that platelets from an AD mouse model damage healthy cortical vessels and induced Aβ accumulation when infused into wildtype OBSC vessels (28). Here we report, for the first time, pathological sprouting angiogenesis in OBSCs from an AD mouse model that is dependent on BACE1 processing of APP. The increased sprouting angiogenesis, including excessive filopodia formation, and increased pericyte coverage, we observe in TgCRND8 OBSCs is much more pronounced than that seen *in vivo* at a similar age, but is consistent with the enhanced vascularisation we observed in both retina and cortex. The OBSC culture method likely stresses the tissue via the slicing injury and/or the isolation from the systemic vasculature resulting in stimulation of sprouting angiogenesis. By allowing visualisation of such processes in both WT and TgCRND8 tissue, the OBSC system appears to unmask a mechanism that underlies the vascular changes in AD models that may otherwise be overlooked.

Use of a model that endogenously expresses mutant APP with all its processing products allows for careful exploration of the effects of Aβ as well as other APP-derived products (36–38). A potential role for Aβ as a regulator of angiogenesis has been previously proposed based on observations in both physiological and pathological conditions. In post mortem human AD brains, it has been shown that increased vascular density in the hippocampus correlates with Aβ load (5). In mouse models of amyloid pathology, immunisation with Aβ peptides resulted in plaque clearance in the brain and restored capillary density to normal levels (39) and inhibition of Aβ oligomerisation/ fibrillization reduced arteriolar Aβ accumulation and tortuosity (6). Application of synthetic Aβ to a number of models has also highlighted its pro-angiogenic role with increased endothelial cell proliferation, capillary bed density and vascular sprouting seen both *in vitro* and *in vivo* (16, 40, 41). Our findings in this study add increasing weight to existing evidence that Aβ or its precursors can stimulate angiogenic processes and we provide a novel OBSC platform to explore the role of endogenously produced Aβ on vascular pathology.

The significance of Aβ here may be as a marker of APP processing or as a cause of pathological angiogenesis. Interestingly, over-production of Aβ in this model is accompanied by reduced expression of the angiogenesis suppressor NOTCH3 and its ligand JAG1, providing a novel mechanistic insight into how amyloid pathology potentially impacts the regulation of angiogenesis in AD. Genetic knockout of *Notch3* has previously been shown to increase retinal vascular density and endothelial tip formation in postnatal mice (42) and silencing NOTCH3 in tumours promotes pathological angiogenesis (43). NOTCH ligand JAG1 has also been implicated in angiogenic processes, with application of Jag1 targeting antisense oligonucleotides potentiating FGF-responsive tube formation and invasion *in vitro* (44). There are multiple potential mechanisms by which *Notch3* and *Jag1* could be downregulated in postnatal TgCRND8 tissue, which we summarise in our working hypothesis (**Fig. 9**).

**Figure 9.**
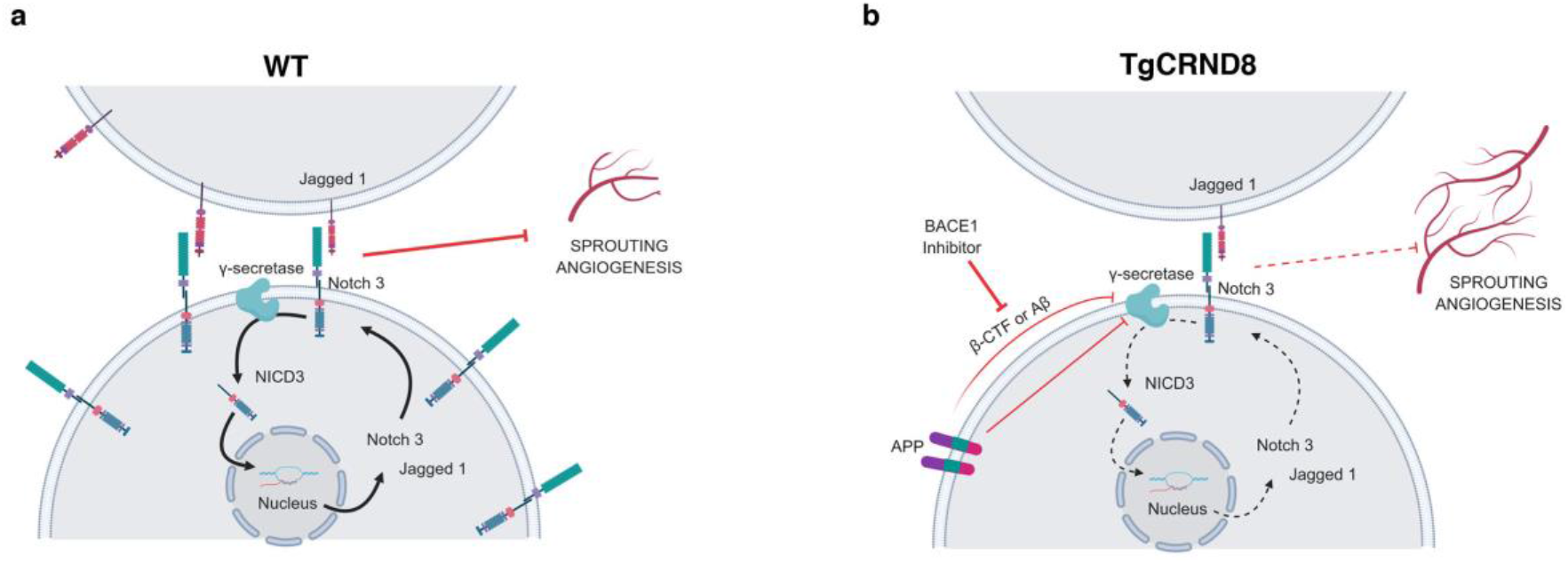
Proposed mechanism for the enhancement of sprouting angiogenesis by BACE1-dependent APP processing. Schematic diagram of our working hypothesis for increased sprouting angiogenesis in TgCRND8 **(b)** compared to WT **(a)** tissue. Increased APP processing by BACE1 in TgCRND8 OBSCs competes with NOTCH3 for γ-secretase or reduces γ-secretase activity, thereby lowering transcriptional signalling through NICD. This reduces *Notch3-Jag1* expression via autoregulatory mechanisms, thereby releasing the inhibitory influence on sprouting angiogenesis.

NOTCH proteins and NOTCH ligands are substrates for the γ-secretase presenilin (45), resulting in the production of NICD which translocates to the nucleus to regulate gene expression (**Fig. 9a**). Cleavage of NOTCH3 by γ-secretase has been found to induce *Notch3* and *Jag1* transcription via autoregulatory mechanisms (46). Previous work has also shown that NOTCH3 activation (by cleavage to NICD3) is prevented by γ-secretase inhibition (47) with γ-secretase treatment found to induce angiogenic sprouting *in vitro* and *in vivo* (48). Interestingly, this effect is mimicked by the application of synthetic monomeric Aβ potentially pointing to an enzymatic feedback inhibition, whereby high levels of Aβ itself also lower the activity of γ-secretase (40). In TgCRND8 tissue (**Fig. 9b**) increased levels of Aβ may act via this mechanism to inhibit the efficacy of γ-secretase, reducing levels of NOTCH3 cleavage and so lowering *Notch3* and *Jag1* transcription, ultimately resulting in increased sprouting angiogenesis. Alternatively, other APP processing products may also have inhibitory effects on γ-secretase. β-CTF, the result of BACE1 cleavage of APP, contains a region (Aβ_17-23_) that has been found to modulate γ-secretase activity by non-competitive inhibition (49) and a similar role has been proposed for the APP intracellular domain (AID) (15). Alternatively, increased expression of APP, or enhanced processing of APP through γ-secretase may directly compete with NOTCH ligands for enzymatic availability. In line with this, there are indications that increased processing of APP through this enzyme competitively downregulates NOTCH1 signalling (50).

One of our important findings is that excessive endothelial filopodia formation and reduced NOTCH3 signalling in TgCRND8 OBSCs can be normalized via application of BACE1 inhibitor (**Fig. 9b**) indicating a likely role for BACE1-dependent APP processing products. BACE1 inhibitors have been shown to inhibit angiogenesis and tumour growth and have been considered as potential cancer therapeutics (51). BACE1 knockout mice show reduced cerebral or retinal vascular density (52) and treatment of zebrafish with BACE1 inhibitors was found to induce similar deficits, with application of Aβ restoring normal vascularisation (53). Whilst we hypothesise that the normalisation of TgCRND8 angiogenesis after BACE1 inhibition is due to reduced levels of BACE-1 dependent APP processing products, we have to consider the possibility that BACE inhibition has a direct effect on angiogenic processes unrelated to APP. Other studies have found that BACE1 directly regulates JAG1 shedding, with BACE1-/- mice showing an increase in *Jag1* levels and downstream NOTCH signalling (54). This, however, seems an unlikely mechanism in our system due to the observation that BACE1 inhibition has no effect on the levels of *Jag1* (**Fig. 7c**) or vascular density (**Fig. 6**) in WT OBSCs.

Further mechanistic studies are needed to better understand the interplay between NOTCH signalling and APP processing mechanisms in AD. Here, we propose that APP overexpression or feedback inhibition from high Aβ/ β-CTF concentration in TgCRND8 OBSCs reduces γ-secretase-dependent cleavage of NOTCH3 (**Fig. 9**). Our results also indicate that there may be a safe level of BACE1 inhibition that can restore physiological levels of angiogenesis, without inducing angiogenic deficits in healthy tissue.

## Methods

### Mice

TgCRND8 mice (55) were maintained on a mixed BL6:sv129 background. TgCRND8 heterozygote males were bred with wild-type background matched females to produce both wild-type and TgCRND8 heterozygote littermates. TgCRND8 mice overexpress human APP with both the Swedish (K670N/M671L) and Indiana (V717F) mutations. Animals were kept on a 12hr:12hr light:dark cycle at a constant temperature of 19°C in a pathogen-free environment. All animal work was approved by the Babraham Institute Animal Welfare and Ethical Review Body and UK Home Office, and carried out in accordance with the Animals (Scientific Procedures) Act, 1986.

### Organotypic brain slice cultures

Cortical organotypic brain slice cultures were made from humanely sacrificed P6-P9 littermates of either sex according to the method of de Simoni et al and our previous work (24, 56). Briefly, after schedule 1, brains were kept on ice in dissection buffer (EBSS supplemented with 25mM HEPES and 1 x Penicillin/Streptomycin). 350μm thick sagittal sections of cortex were cut using a Leica VT1000S vibratome and slices collected using a modified sterile 3ml Pasteur pipette. On average 6 cortical slices were collected per pup and stored in dissection buffer on ice until plating. For long term culture, slices were transferred, in a class II hood, onto sterile Millicell^®^ membrane inserts (Millipore PICM0RG50) in 35mm culture dishes (Nunc). 3 cortical slices from the same pup were plated to a single membrane, with two dishes made per animal. Inserts were maintained in 1ml of maintenance medium according to the following recipe: 50% MEM with Glutamax-1 (Life Technologies 42360-024), 25% Heat-inactivated horse serum (Life Technologies: 26050-070), 23% EBSS (Life Technologies: 24010-043), 0.65% D-Glucose (Sigma:G8270), 2% Penicillin/Streptomycin (Life Technologies 15140-122) and 6 units/ml Nystatin (Sigma N1638). Membrane inserts were handled with sterile forceps and the medium was changed 100% 4 hours after plating and at 4 *div*. OBSCs were maintained at 37°C, 5% CO_2_ and high humidity for 7 *div*. For BACE1 inhibition experiments, 1 culture per pup was treated with 5μM BACE1 inhibitor LY2886721 (Selleckchem S2156) and compared with a DMSO treated control from the same animal. Cultures were treated for the entire 7 days *in vitro*.

### Immunohistochemistry

OBSCs were fixed for 20 minutes in 4% PFA in phosphate buffered saline (PBS), washed 3 times in PBS, then blocked for 1 hour in PBS supplemented with 0.5% Triton X-100 and 3% normal Goat Serum (Sigma G9023). Slices were incubated at 4°C in primary antibody (diluted in blocking solution) overnight. In order to detect PDGFR-β, heat-mediated antigen retrieval was performed in 10 mM citrate buffer (pH 6.0) for 40 min at 80°C prior to primary antibody incubation. Slices were washed a further 3 times in 0.5% Triton-X100 in PBS (PBS-TX) then incubated with secondary antibodies (Life Technologies and Jackson) (1:500 dilution in blocking solution for 2 hours at 4°C). Three final PBS-TX washes were conducted before slices were mounted on slides and images captured using a Leica Confocal Microscope. Primary antibodies used: rabbit anti-PDGFR-β (28E1) (1:200, Cell Signalling), rat anti-PECAM-1 (1:400, BD), rabbit anti-laminin (1:200, Sigma) secondary staining was conducted using species-specific fluorophore-conjugated (Streptavidin Alexa 488, Molecular Probes; Cy3 or Cy5, Jackson,) or biotin-conjugated secondary antibodies (Jackson). DAPI (1μg/mL, Sigma) was used to counterstain nuclei.

### Microscopy and Image Analysis

Images were captured via an epifluorescence microscopy system (Leica DM6000B) or using confocal microscopes (Leica, Zeiss LSM780). Figures were composed using Photoshop CS5 software.

#### Quantification of pericyte number and coverage

To quantify the number of PDGFRβ-positive pericytes, cells were counted using NIH Image J Cell Counter tool. A maximum projection of fifteen-micrometre z-stacks was acquired from cortex slices freshly derived from WT and TgCRND8. Three pictures were taken at 40X for each slice. The areas of PDGFR-β positive pericytes and PECAM-1-positive blood vessels from were separately subjected to threshold processing and the respective signals for each image was calculated using NIH Image J Area Measurement tool. Pericyte coverage was determined as a percentage (%) of PDGFRβ--positive pericyte area covering PECAM-1-positive capillary surface area per field (ROI) 733×733 μm. Three slices per animal were analysed.

#### Capillary density, length and filopodia quantification

For PECAM-1-postive capillary area, sections were analysed with Leica confocal microscope. Three pictures were taken at 20X for each section. ROI size of 733×733 μm for confocal images were utilized. The area covered by PECAM-1-positive capillaries was analysed using the NIH ImageJ area measurement tool where pictures were subjected to threshold processing to produce a binary image. The area of PECAM-1 coverage was expressed as percentage of the total area, 3 slices per WT and TgCRND8 pups (n= 4-5 for each genotype) were analysed. The filopodia of vascular sprouts were analysed using z-stacked PECAM-1 positive blood vessels.

### Protein extraction and Western Blot

OBSCs were scraped off the membrane insert using a scalpel and transferred to 2 x Laemelli buffer supplemented with 10% 2-mercaptethanol (50μL per 3 slices). Samples were boiled for 10 minutes then frozen at −20°C until use. Ten micrograms of protein were separated on a 10% SDS polyacrylamide gel then transferred onto PVDF membranes. Membranes underwent blocking (20 mM Tris, 136 mM NaCl, pH 7.6, 0.1% Tween 20, 5% nonfat dry milk) before incubation with primary antibody anti-NOTCH1/NICD1 (1:750, Abcam) NOTCH3/NICD3 (1:1000, Abcam) PDGFR-β (28E1) (1:1000, Cell Signaling) overnight at 4°C. overnight at 4 °C. Signals were obtained by binding of a secondary anti-rabbit horse radish peroxidase (HRP) linked antibody (1:15000, Sigma-Aldrich) and visualized by exposing the membrane to a charge-coupled device camera (LAS1000, Fujifilm, Tokyo, Japan) using a chemiluminescence kit (Merck Millipore, Billerica, MA, USA). Membranes were stripped and reprobed for β-actin (diluted 1:50 000, Sigma-Aldrich). After densitometric analysis using Image J software, protein levels were calculated as percentage of β-actin expression.

### Aβ ELISA

Culture medium from 7 *div* OBSCs was assayed for human Aβ_1-40_ and Aβ_1-42_ using commercially available ELISA kits (Life Technologies: KHB3441 and KHB3481). Briefly, medium was incubated with Aβ detection antibody for 3 hours, washed, and then incubated with an HRP-conjugated antibody for 30 minutes. After another wash step, stabilised chromogen was added for 30 minutes before the adding an acid-based stop solution. Absorbance was read at 450nm using a PheraStar ELISA plate reader with the standard curve calculated using a 4-parameter fit. Concentration of Aβ in the medium is expressed as pg/ml. Levels of Aβ were compared between BACE1 treated and DMSO treated TgCRND8 cultures.

### Quantification of gene expression by qPCR

Slice cultures were scraped off the membrane and RNA extracted using an RNeasy mini kit (Qiagen). Briefly, 3 slices were homogenised in 350μL lysis buffer RLT supplemented with 1% 2-mercaptethanol. 350μL 70% ethanol (in nuclease free water) was then added and samples transferred to RNeasy RNA collection columns. After several wash steps described in the kit protocol, the RNA was eluted in 20μL nuclease free water, measured and quality tested using a Nanodrop^®^ and frozen at −80°C until use. cDNA synthesis was performed using Script^™^ cDNA Synthesis Kit (Bio-Rad). cDNA was analysed using real-time PCR SsoAdvanced^™^ SYBR^®^ Green Supermix from Bio-Rad and run on a Bio-Rad CFX96 real-time quantitative PCR (qPCR) system. Gene expression was normalized to the housekeeping gene GAPDH. Melt curve analyses were performed to ensure the specificity of qPCR product. Primer sequences will be provided on request. Values are presented as mean ± SD of three independent experiments, and within each experiment, triplicate samples were assessed.

### Statistical Analysis

Statistical analysis was conducted using Graph Pad Prism. Data are expressed as mean ± SD. Two group comparison was performed by using Student’s t-test and multiple group comparison by one-way or two-way ANOVA followed by Tukey post hoc test. For Western blot, after normalisation to the actin signal, the ipsilateral expression of each junction protein was compared with the using a two-way ANOVA, followed by Tukey post hoc test. Significance was set at p<0.05.

## Data availability

The datasets generated and analyzed in this study are available from the corresponding author on request.

## Author Contributions

C.S.D performed research, analysed the data and wrote the manuscript; K.R performed research and analysed the data; O.S. performed research; M.P.C. designed research and wrote the manuscript; I.O. designed the research, analysed the data and wrote the manuscript.

## References

1. Jefferies WA, et al. (2013) Adjusting the compass: new insights into the role of angiogenesis in Alzheimer’s disease. Alzheimers Res Ther 5(6):64.

2. Giuliani A, et al. (2019) Age-Related Changes of the Neurovascular Unit in the Cerebral Cortex of Alzheimer Disease Mouse Models: A Neuroanatomical and Molecular Study. J Neuropathol Exp Neurol 78(2):101–112.

3. Yamazaki Y, et al. (2019) Selective loss of cortical endothelial tight junction proteins during Alzheimer’s disease progression. Brain J Neurol. doi:10.1093/brain/awz011.

4. Biron KE, Dickstein DL, Gopaul R, Jefferies WA (2011) Amyloid triggers extensive cerebral angiogenesis causing blood brain barrier permeability and hypervascularity in Alzheimer’s disease. PloS One 6(8):e23789.

5. Desai BS, Schneider JA, Li J-L, Carvey PM, Hendey B (2009) Evidence of angiogenic vessels in Alzheimer’s disease. J Neural Transm Vienna Austria 1996 116(5):587–597.

6. Dorr A, et al. (2012) Amyloid-β-dependent compromise of microvascular structure and function in a model of Alzheimer’s disease. Brain 135(10):3039–3050.

7. Lai AY, et al. (2015) Venular degeneration leads to vascular dysfunction in a transgenic model of Alzheimer’s disease. Brain 138(4):1046–1058.

8. Meyer EP, Ulmann-Schuler A, Staufenbiel M, Krucker T (2008) Altered morphology and 3D architecture of brain vasculature in a mouse model for Alzheimer’s disease. Proc Natl Acad Sci U S A 105(9):3587–3592.

9. Berisha F, Feke GT, Trempe CL, McMeel JW, Schepens CL (2007) Retinal abnormalities in early Alzheimer’s disease. Invest Ophthalmol Vis Sci 48(5):2285–2289.

10. Hardy J (2002) The Amyloid Hypothesis of Alzheimer’s Disease: Progress and Problems on the road to Therapeutics. Science 297(5580):353–356.

11. Terry RD, et al. (1991) Physical basis of cognitive alterations in Alzheimer’s disease: synapse loss is the major correlate of cognitive impairment. Ann Neurol 30.

12. Potente M, Gerhardt H, Carmeliet P (2011) Basic and therapeutic aspects of angiogenesis. Cell 146(6):873–887.

13. Phng L-K, Gerhardt H (2009) Angiogenesis: a team effort coordinated by notch. Dev Cell 16(2):196–208.

14. Hartmann D, Tournoy J, Saftig P, Annaert W, De Strooper B (2001) Implication of APP secretases in notch signaling. J Mol Neurosci MN 17(2):171–181.

15. Roncarati R, et al. (2002) The gamma-secretase-generated intracellular domain of beta-amyloid precursor protein binds Numb and inhibits Notch signaling. Proc Natl Acad Sci U S A 99(10):7102–7107.

16. Boscolo E, et al. (2007) β amyloid angiogenic activity in vitro and in vivo. Int J Mol Med 19(4):581–587.

17. Ethell DW (2010) An Amyloid-Notch Hypothesis for Alzheimer’s Disease. The Neuroscientist 16(6):614–617.

18. Hellström M, Kalén M, Lindahl P, Abramsson A, Betsholtz C (1999) Role of PDGF-B and PDGFR-beta in recruitment of vascular smooth muscle cells and pericytes during embryonic blood vessel formation in the mouse. Dev Camb Engl 126(14):3047–3055.

19. Lindblom P, et al. (2003) Endothelial PDGF-B retention is required for proper investment of pericytes in the microvessel wall. Genes Dev 17(15):1835–1840.

20. Hutter-Schmid B, Kniewallner KM, Humpel C (2015) Organotypic brain slice cultures as a model to study angiogenesis of brain vessels. Front Cell Dev Biol 3. doi:10.3389/fcell.2015.00052.

21. Moser KV, Schmidt-Kastner R, Hinterhuber H, Humpel C (2003) Brain capillaries and cholinergic neurons persist in organotypic brain slices in the absence of blood flow. Eur J Neurosci 18(1):85–94.

22. Moser KV, Reindl M, Blasig I, Humpel C (2004) Brain capillary endothelial cells proliferate in response to NGF, express NGF receptors and secrete NGF after inflammation. Brain Res 1017(1):53–60.

23. Croft CL, Noble W (2018) Preparation of organotypic brain slice cultures for the study of Alzheimer’s disease. F1000Research 7. doi:10.12688/f1000research.14500.2.

24. Harwell CS, Coleman MP (2016) Synaptophysin depletion and intraneuronal Aβ in organotypic hippocampal slice cultures from huAPP transgenic mice. Mol Neurodegener 11:44.

25. Holopainen IE (2005) Organotypic Hippocampal Slice Cultures: A Model System to Study Basic Cellular and Molecular Mechanisms of Neuronal Cell Death, Neuroprotection, and Synaptic Plasticity. Neurochem Res 30(12):1521–1528.

26. Humpel C (2015) Organotypic vibrosections from whole brain adult Alzheimer mice (overexpressing amyloid-precursor-protein with the Swedish-Dutch-Iowa mutations) as a model to study clearance of beta-amyloid plaques. Front Aging Neurosci 7:47.

27. Novotny R, et al. (2016) Conversion of Synthetic Aβ to In Vivo Active Seeds and Amyloid Plaque Formation in a Hippocampal Slice Culture Model. J Neurosci 36(18):5084–5093.

28. Kniewallner KM, Foidl BM, Humpel C (2018) Platelets isolated from an Alzheimer mouse damage healthy cortical vessels and cause inflammation in an organotypic ex vivo brain slice model. Sci Rep 8. doi:10.1038/s41598-018-33768-2.

29. Chung AS, Ferrara N (2011) Developmental and pathological angiogenesis. Annu Rev Cell Dev Biol 27:563–584.

30. Ozerdem U, Stallcup WB (2003) Early contribution of pericytes to angiogenic sprouting and tube formation. Angiogenesis 6(3):241–249.

31. Cheung CY-L, et al. (2014) Microvascular network alterations in the retina of patients with Alzheimer’s disease. Alzheimers Dement J Alzheimers Assoc 10(2):135–142.

32. Farkas E, Luiten PG (2001) Cerebral microvascular pathology in aging and Alzheimer’s disease. Prog Neurobiol 64(6):575–611.

33. Ullrich C, Humpel C (2009) The Pro-Apoptotic Substance Thapsigargin Selectively Stimulates Re-Growth of Brain Capillaries. Curr Neurovasc Res 6(3):171–180.

34. Chip S, Zhu X, Kapfhammer JP (2014) The Analysis of Neurovascular Remodeling in Entorhino-hippocampal Organotypic Slice Cultures. J Vis Exp (92). doi:10.3791/52023.

35. Daschil N, et al. (2015) L-type calcium channel blockers and substance P induce angiogenesis of cortical vessels associated with beta-amyloid plaques in an Alzheimer mouse model. Neurobiol Aging 36(3):1333.

36. Moore S, et al. (2015) APP Metabolism Regulates Tau Proteostasis in Human Cerebral Cortex Neurons. Cell Rep 11(5):689–696.

37. Walsh DM, Klyubin I, Fadeeva JV, Rowan MJ, Selkoe DJ (2002) Amyloid-beta oligomers: their production, toxicity and therapeutic inhibition. Biochem Soc Trans 30.

38. Willem M, et al. (2015) η-Secretase processing of APP inhibits neuronal activity in the hippocampus. Nature 526(7573):443–447.

39. Biron KE, Dickstein DL, Gopaul R, Fenninger F, Jefferies WA (2013) Cessation of Neoangiogenesis in Alzheimer’s Disease Follows Amyloid-beta Immunization. Sci Rep 3:1354.

40. Cameron DJ, et al. (2012) Alzheimer’s-Related Peptide Amyloid-β Plays a Conserved Role in Angiogenesis. PLOS ONE 7(7):e39598.

41. Cunvong K, Huffmire D, Ethell DW, Cameron DJ (2013) Amyloid-β Increases Capillary Bed Density in the Adult Zebrafish Retina. Invest Ophthalmol Vis Sci 54(2):1516–1521.

42. Kofler NM, Cuervo H, Uh MK, Murtomäki A, Kitajewski J (2015) Combined deficiency of Notch1 and Notch3 causes pericyte dysfunction, models CADASIL and results in arteriovenous malformations. Sci Rep 5(1). doi:10.1038/srep16449.

43. Lin S, et al. (2017) Non-canonical NOTCH3 signalling limits tumour angiogenesis. Nat Commun doi:10.1038/ncomms16074.

44. Zimrin AB, et al. (1996) An Antisense Oligonucleotide to the Notch Ligand Jagged Enhances Fibroblast Growth Factor-induced Angiogenesis in Vitro. J Biol Chem 271(51):32499–32502.

45. Groot AJ, et al. (2014) Regulated Proteolysis of NOTCH2 and NOTCH3 Receptors by ADAM10 and Presenilins. Mol Cell Biol 34(15):2822–2832.

46. Liu H, Kennard S, Lilly B (2009) NOTCH3 expression is induced in mural cells through an autoregulatory loop that requires endothelial-expressed JAGGED1. Circ Res 104(4):466–475.

47. Konishi J, et al. (2007) Gamma-secretase inhibitor prevents Notch3 activation and reduces proliferation in human lung cancers. Cancer Res 67(17):8051–8057.

48. Kalén M, et al. (2011) Gamma-Secretase Inhibitor Treatment Promotes VEGF-A-Driven Blood Vessel Growth and Vascular Leakage but Disrupts Neovascular Perfusion. PLOS ONE 6(4):e18709.

49. Zhang Y, Xu H (2010) Substrate check of γ-secretase. Nat Struct Mol Biol 17(2):140–141.

50. Berezovska O, et al. (2001) Notch1 and Amyloid Precursor Protein Are Competitive Substrates for Presenilin1-dependent γ-Secretase Cleavage. J Biol Chem 276(32):30018–30023.

51. Paris D, et al. (2005) Inhibition of angiogenesis and tumor growth by β and γ-secretase inhibitors. Eur J Pharmacol 514(1):1–15.

52. Cai J, et al. (2012) β-Secretase (BACE1) inhibition causes retinal pathology by vascular dysregulation and accumulation of age pigment. EMBO Mol Med 4(9):980–991.

53. Luna S, Cameron DJ, Ethell DW (2013) Amyloid-β and APP Deficiencies Cause Severe Cerebrovascular Defects: Important Work for an Old Villain. PLoS ONE 8(9). doi:10.1371/journal.pone.0075052.

54. Hu X, He W, Luo X, Tsubota KE, Yan R (2013) BACE1 regulates hippocampal astrogenesis via the Jagged1-Notch pathway. Cell Rep 4(1):40–49.

55. Chishti MA, et al. (2001) Early-onset Amyloid Deposition and Cognitive Deficits in Transgenic Mice Expressing a Double Mutant Form of Amyloid Precursor Protein 695. J Biol Chem 276(24):21562–21570.

56. De Simoni A, MY Yu L (2006) Preparation of organotypic hippocampal slice cultures: interface method. Nat Protoc 1(3):1439–1445.

